# The licorice metabolite enoxolone attenuates *Clostridioides difficile* pathophysiology by corrupting its metabolic and toxin production networks

**DOI:** 10.1101/2022.04.20.488993

**Authors:** Ravi K. R. Marreddy, Jonathan Picker, Gregory A. Phelps, Reid Powell, Philip T. Cherian, John J. Bowling, Clifford C. Stephan, Richard E. Lee, Julian G. Hurdle

## Abstract

Toxins TcdA and TcdB are the main virulence factors of *Clostridioides difficile*, a leading cause of hospital-acquired diarrhea. We investigated the therapeutic potential of inhibiting the biosynthesis of TcdA and TcdB. Accordingly, screening of structurally diverse phytochemicals with medicinal properties identified 18β-glycyrrhetinic acid (enoxolone) as an inhibitor of TcdA and TcdB biosynthesis. Enoxolone also inhibited sporulation. In a CDI colitis model, enoxolone when combined with vancomycin protected mice from becoming moribund and the combination was more effective than vancomycin alone, a standard of care antibiotic for CDI. While enoxolone alone reduced the *in vivo* load of toxins, the monotherapy did not protect mice from CDI. Affinity based proteomics identified ATP synthase subunit alpha (AtpA) and adenine deaminase (Ade) as possible molecular targets for enoxolone. Silencing of mRNA for Ade and AtpA also reduced toxin biosynthesis, while molecular interaction analysis showed that enoxolone directly bound to Ade. Ade converts adenine to hypoxanthine as an early step in the purine salvage pathway. Metabolomics revealed enoxolone caused cells to accumulate adenosine and deplete hypoxanthine and ATP. Accordingly, supplementation with hypoxanthine partly restored toxin production. Enoxolone also impacted phosphate metabolism by reducing the amounts of cellular phosphate. Thus, supplementation with triethyl phosphate as a source of phosphate also partly restored toxin production. When hypoxanthine and triethyl phosphate were combined, toxin production was fully restored in the presence of enoxolone. Taken together, studies with enoxolone revealed metabolic pathways that affect *C. difficile* toxin production and could represent potential anti-virulence drug targets.

**IMPORTANCE:** *Clostridioides difficile*, a leading cause of hospital-acquired diarrhea, produces two co-regulated toxins (TcdA and TcdB) that are the focus of most anti-virulence discovery efforts for *C. difficile* infection (CDI). Exploration of an alternate anti-virulence strategy led to the discovery that the licorice metabolite enoxolone inhibits *C. difficile* virulence by blocking the cellular biosynthesis of TcdA and TcdB. Blockage of toxin production by enoxolone was associated with multiple effects on cells, including inhibiting adenine deaminase and ATP synthase leading to disruption of purine biosynthesis and phosphate metabolism. In mice infected with *C. difficile*, the efficacy of enoxolone in combination with vancomycin was superior to vancomycin alone. These findings contribute to establishing toxin biosynthesis inhibition as a newer therapeutic concept for CDI.

## INTRODUCTION

*Clostridioides difficile* infection (CDI) is a leading cause of hospital-acquired diarrhea, causing 30,600 and 20,500 in-hospital deaths in the U.S. from 462,100 and 476,400 cases in 2011 and 2017, respectively (1). Various broad-spectrum antibiotics are leading risk factors for CDI, as they promote dysbiosis and *C. difficile* colonization (2). While metronidazole and vancomycin have been the main antibiotic therapies prescribed for CDI, 20% or more patients experience recurrence following therapy with these agents (3). These two antibiotics also inhibit the growth of important gut flora and are thought to further disturb the microbiota. Therefore, there is a medical need for novel CDI therapeutics that are more targeted to *C. difficile* pathophysiology.

*C. difficile* produces two glycosyltransferase toxins (TcdA and TcdB), in the late-log and stationary phases of growth, which are responsible for CDI symptoms (4). Thus, significant research attention has been given to developing biotherapeutics and small molecules that inactivate *C. difficile* toxins (5–7). The leading example is bezlotoxumab, a monoclonal antibody to TcdB, which is combined with standard-of-care antibiotics to treat recurrent CDI (6). Nonetheless, there is a paucity of discovery-oriented research to identify inhibitors that block cellular synthesis of TcdA and TcdB. Biosynthesis of TcdA and TcdB is regulated by various sigma factors, environmental and nutritional changes (8). For example, during nutrient excess and high intracellular concentrations of GTP, the nutritional regulator CodY binds to the *tcdR* promoter to suppress *tcdA* and *tcdB* transcription (9, 10); glucose, and other metabolizable carbohydrates, impedes toxin production through catabolite repression, whereby the catabolite control protein A (CcpA) binds to the promoter of the alternative sigma factor *tcdR*, whose gene product activates *tcdA* and *tcdB* (11–13); and activation of Stickland metabolism of proline, which is thought to suppresses toxin biosynthesis (14). While these and other pathways affect toxin biosynthesis, there have been limited systematic efforts to identifying inhibitors of toxin production (15).

To identify potential toxin biosynthesis inhibitors, we focused on a small panel of naturally occurring phytochemicals from different chemical classes (**Table S1**). The potential for medicinal and food-based phytochemicals as treatments for CDI is underscored by berberine and curcumin that are reported to be efficacious in murine models of CDI; their efficacies likely result from inhibition of spore germination and toxin production, respectively (16, 17). We also reported that naturally occurring flavonoids have weak antibacterial activity against *C. difficile* but did not affect toxin biosynthesis unless chemically modified to be potent antibacterials (18). However, the fact that plant-derived compounds often display weak antibacterial activities (19), imply that they could be promising chemical starting points for inhibitors of TcdA and TcdB biosynthesis. We therefore explored different classes of phytochemicals to discover that the pentacyclic triterpenoid 18β-glycyrrhetinic acid (enoxolone) blocks cellular production of TcdA and TcdB. This occurred through a complex mode of action that reconfigured *C. difficile* purine, ATP and phosphate metabolism in a manner that was unfavorable for toxin production.

## RESULTS

### Screening for toxin biosynthesis inhibitors for *C. difficile*

We assembled a small panel (n=57) of phytochemicals/metabolites (**Table S1**) from differing structural classes to identify molecules that inhibit toxin production in *C. difficile* R20291, without substantially preventing growth. **Table S1** describes these phytochemicals, their sources, and medicinal properties, where available. To identify inhibitors, we tested molecules at 100 μM and triaged those that inhibited growth by ≥50%. After triaging growth inhibitors, toxins were quantified in the cultures exposed to the remaining compounds. This was done using a cytopathic assay that was based on TcdB-induced rounding of Vero cells (20) and quantifying toxins from automated morphometric analysis of cell shape and surface area covered in 384-well plates. Similar automated assays have been reported for *C. difficile* TcdB (21). Presumptive compound hits were then confirmed by ELISA to quantify the amounts of TcdA and TcdB in the supernatants. Results showed that of the 57 compounds, 6 were eliminated as they inhibited growth by >60% (**Figure 1A**). To select hits from the remaining 51 compounds, we analyzed the cytotoxic effects of culture supernatants on Vero cells. The supernatants obtained from *C. difficile* cells treated with DMSO caused 90.86 ± 0.69% of cell rounding, while cell rounding by culture supernatants from glucose (1% w/v) control treated cells was 20.31 ± 4.89% (**Figure 1B**). Upon comparison, four hits were identified that attenuated *C. difficile* cytopathy, in a manner comparable to glucose: enoxolone (15.02 ± 0.87%), parthenolide (16.21 ± 1.15%), kahweol (20.74 ± 8.55%) and tannic acid (14.57 ± 2.71%). ELISA performed on the supernatants confirmed that TcdA and TcdB concentrations were low (**Figure 1C**); the compounds decreased TcdA by ≥70% and TcdB by ≥50% and were comparable to glucose. The comparator vancomycin did not inhibit toxin production when used at a growth permissive concentration (0.5 μM). Next, when dose-responses were determined using the cell rounding assay, kahweol and tannic acid were found to have high EC_50_ values (>80 µM), when compared to parthenolide (EC_50_=27.93 µM) and enoxolone (7.77 µM) (**Figure 1D**). This data suggested that enoxolone and parthenolide are inhibitors of toxin production in *C. difficile*.

**Figure-1.**
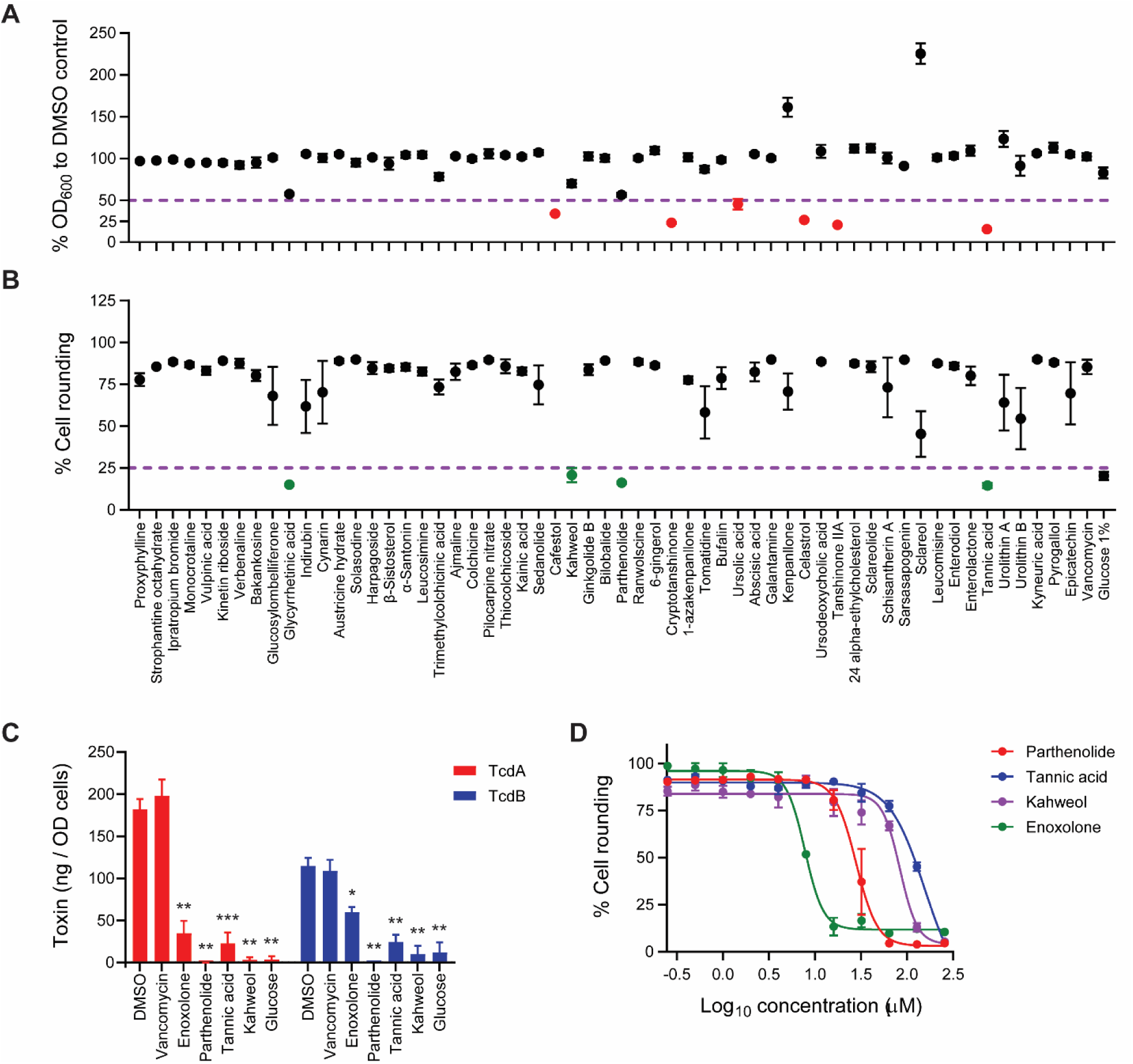
Screening for inhibitors of *C. difficile* toxin biosynthesis. R20291 in BHI in 96-well plates was exposed to 100 µM of phytochemicals for 24 h. Three biological replicates were analyzed for: (**A**) effects on growth (OD_600_nm); growth inhibitors are indicated in red and were triaged by comparison with the DMSO control. (**B**) Toxins were quantified by cytopathic cell rounding; inhibitors of toxin synthesis are indicated in green (n = 4 biological replicates). Controls were vancomycin (0.5 µM) and glucose 1% (w/v). The purple line indicates cutoff criteria for compound selection. (**C**) Toxins (TcdA and TcdB) in culture supernatants were quantified by ELISA; three biological replicates were used and shown as means ± standard error of mean (SEM); statistical significance was assessed by unpaired t-test with Welch’s correction *, P<0.05; **, P<0.01; ***, P<0.001. (**D**) Dose-response curves of toxin biosynthesis inhibitors parthenolide, tannic acid, kahweol, and enoxolone. Toxins, namely TcdB, were quantified by cell rounding against Vero cells and used four biological replicates (data is shown as mean ± SEM).

### Characterization of the anti-virulence and antibacterial properties of enoxolone

Enoxolone is a pentacyclic triterpenoid aglycone metabolite of glycyrrhizin that is found in licorice. It exhibits a range of pharmacological properties including antitumor, anti-inflammatory, and anti-viral activities (22). We decided to focus on enoxolone due its greater potency, chemical tractability, and reports of related pentacyclic triterpenoids that show anti-virulence against various bacteria (23). We first confirmed EC_50_s by ELISA and cell rounding against various ribotype 027 strains. The EC_50_s for inhibition of TcdA and TcdB production (**Figures 2A, S1A and S1B**) were similar to the EC_50_s observed in the cell rounding assay. Based on findings against ribotype 027 strains, we selected 16 µM as the effective concentration for further experimentation. Upon exposure to enoxolone for 9 h, measurement of *tcdA* and *tcdB* transcripts revealed a dose-dependent effect, where 8 µM (0.5 x EC_50_ for toxin biosynthesis inhibition) did not have a significant effect, while 16 µM and 32 µM (1 x and 2 x EC_50_ for toxin biosynthesis inhibition, respectively) reduced transcription as follows: (*tcdA* [2.17 ± 0.49 and 10.1 ± 4.2 –fold]) and *tcdB* ([1.8 ± 0.3 and 2.9 ± 0.36 –fold]) respectively (**Figure 2B**). Because *tcdA* is more highly expressed than *tcdB* (13), and its expression is indicative for *tcdB*, we subsequently measured TcdA in cells as a readout for toxin production. We next analyzed the activity of enoxolone against major CDI-associated ribotypes (PCR ribotypes 014, 020, 078 and 106). Enoxolone (32 µM) reduced toxin production by >80% in most strains, except for ribotype-020 where the reduction was ∼50% (**Figure S1C and S1D**). These findings indicate that enoxolone is active against different CDI ribotypes.

**Figure-2.**
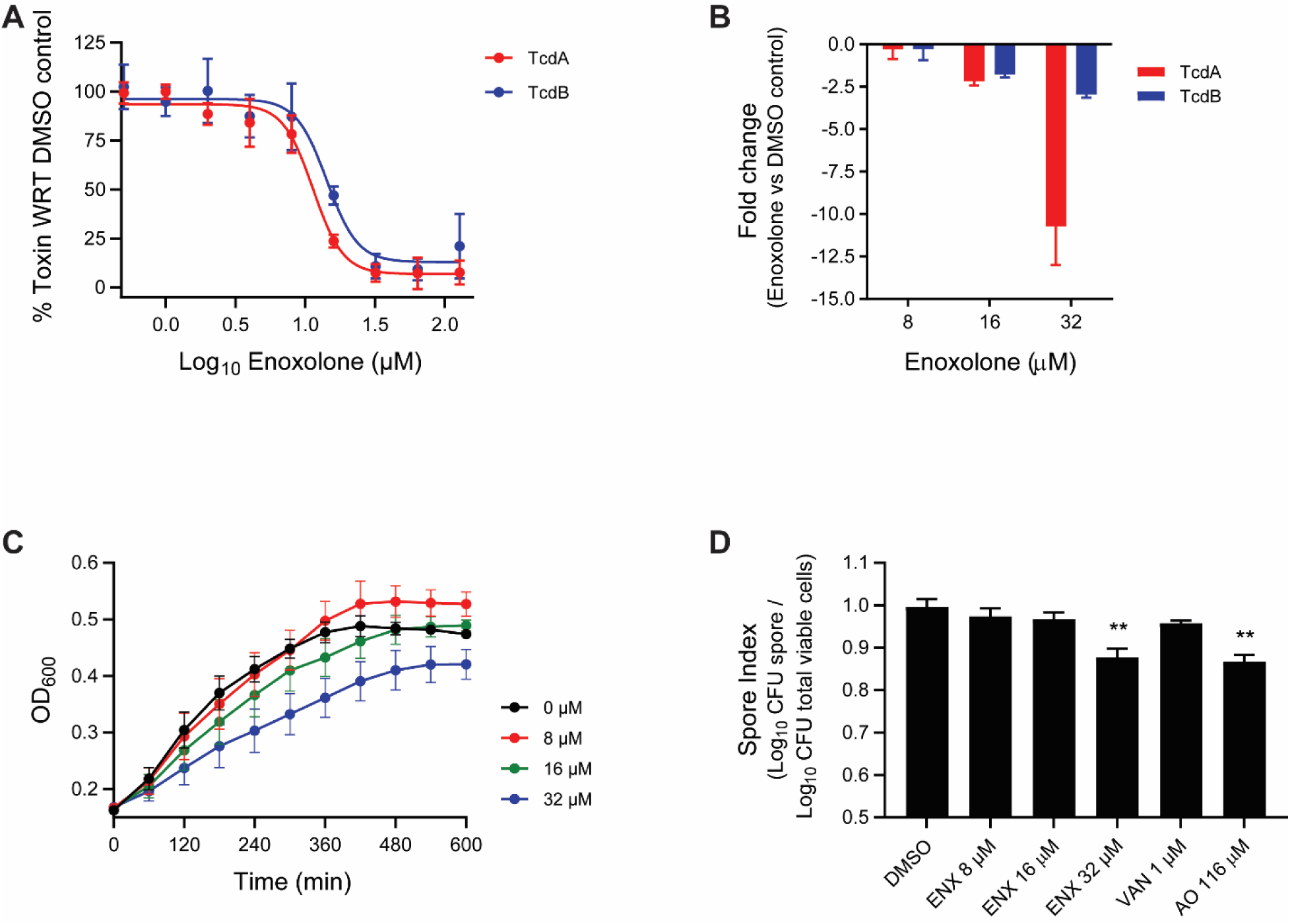
Characterization of enoxolone (ENX) activity against *C. difficile*. **(A)** Dose response of enoxolone against R20291. Exponential cultures (OD_600_nm ≈ 0.3) were exposed to DMSO or 2-fold increasing concentrations of enoxolone (n = 3 biological replicates). After 24 h, TcdA and TcdB were quantified by ELISA. **(B)** Effect of enoxolone on mRNA levels of *tcdA* and *tcdB*, as determined by RT-qPCR. Cultures (OD_600_ ≈ 0.3) were exposed to enoxolone, and mRNA analyzed after 9 h; the fold change was calculated relative to the DMSO control. **(C)** Effect of enoxolone on growth of R20291. Exponential cells (OD_600_nm ≈ 0.2; n=4 biological replicates) were treated with DMSO or enoxolone at ½ x, 1 x and 2 x EC_50_ of 8 μM. (**D**) Effect of enoxolone on sporulation. Exponential cells (OD_600_nm ≈ 0.3; n=4 biological replicates) were exposed to compound or DMSO for 5 days; total viable counts and spores were then enumerated; enoxolone was used at ½ x, 1 x and 2 x EC_50_ of 8 μM. Controls were vancomycin (1 µM) and acridine orange (AO=116 µM).

Screening of enoxolone at 100 µM reduced cell biomass (i.e., OD_600_ nm) by ∼40% at 24 h endpoint. Thus, to better assess the effects of enoxolone, we analyzed the kinetic growth of R20291 in various concentrations of the compound. This revealed that enoxolone at 32 µM delayed growth but had a modest effect on growth at the EC_50_ (16 µM) for inhibition of toxin production (**Figure 2C**). Against various gut flora species, enoxolone did not inhibit the growth of all tested organisms at concentrations below 128 µM. In contrast, vancomycin inhibited the growth of these microorganisms with MICs of 0.5 to 16 µM, except for *Bacteroides* spp. (MIC≥32 µM) (**Table 1**).

**Table-1.** Antimicrobial activity of enoxolone (ENX) and vancomycin (VAN), in BHI broth, against various gut microbial species.

| Bacteria | Strain | MIC (μM) | |
| --- | --- | --- | --- |
|  |  | ENX | VAN |
| *Actinomyces viscosus* | HM238 | >128 | 0.5 - 1 |
| *Bacteroides fragilis* | HM20 | >128 | 8 |
| *Bacteroides eggerthii* | HM210 | >128 | 8 |
| *Bacteroides ovatus* | HM222 | >128 | $\geq 32$ |
| *Bacteroides Sp.* | HM18 | >128 | $\geq 32$ |
| *Bacteroides Sp.* | HM19 | >128 | $\geq 32$ |
| *Bacteroides Sp.* | HM23 | >128 | $\geq 32$ |
| *Bacteroides Sp.* | HM28 | >128 | $\geq 32$ |
| *Fusobacterium nucleatum* | HM260 | >128 | 16 |
| *Lactobacillus Johnsonii* | HM643 | >128 | 0.5 - 1 |
| *Lactobacillus crispatus* | HM421 | >128 | 0.5 – 1 |
| *Porphyromonas uenonis* | HM130 | 64 - >128 | 1 - 4 |

Because regulatory mechanisms for toxin production can intersect with those for sporulation in *C. difficile* (8), we tested the effect of enoxolone on sporulation. Cultures were treated with compounds for 5 days before measuring total viable counts and spores (24). This revealed that enoxolone only inhibited sporulation at 32 µM (**Figures 2D**), which was comparable to the positive control acridine orange (116 µM). In contrast, the comparator vancomycin (at its MIC of 1 μM), did not inhibit sporulation (**Figure 2D**).

### Efficacy of enoxolone in a CDI colitis mouse model

To test the efficacy of enoxolone we modified a reported CDI colitis model (25), by using a lower concentration of dextran sodium sulfate (DSS; 1.0% w/v instead of 3% w/v) and administering antibiotics and DSS for 5 days instead of 3 days (**Figure 3A**). These modifications prevented colitis-related morbidity (25) and facilitated the reproducible development of CDI. We conducted two independent experiments in which male and female mice were infected with R20291 spores and treated twice daily with either vehicle, enoxolone (50 mg/kg), vancomycin (20 mg/kg) or combinations of vancomycin and enoxolone. Treatment with enoxolone began 2 hours after infection, while treatment with vancomycin was started 24 hours after infection. We observed that the overall morbidities and mortalities of mice treated with enoxolone alone were similar to those given the vehicle (**Figure 3B** to **3D**). Compared to the vehicle group, mice that were only given enoxolone experienced greater initial weight loss (**Figure 3C**). Weight loss was also evident in the enoxolone/vancomycin combination group, when compared to mice that were only given vancomycin (**Figure 3D**). We speculate that the observed increased weight loss was a side effect of enoxolone, as the compound is reported to cause hypertension and an analog was shown to suppress food intake (26). Analysis of spore and toxin loads revealed there was no significant difference in spore bioburdens (**Figure 3E**), but the fecal toxin titers of enoxolone treated mice were ∼1 log less than mice given the vehicle (**Figure 3G**). However, the fact that mice became moribund when given the monotherapy suggests that the toxin titers were not sufficiently decreased to prevent CDI symptoms and that enoxolone may be better exploited as an adjuvant to antibiotic therapy.

**Figure-3.**
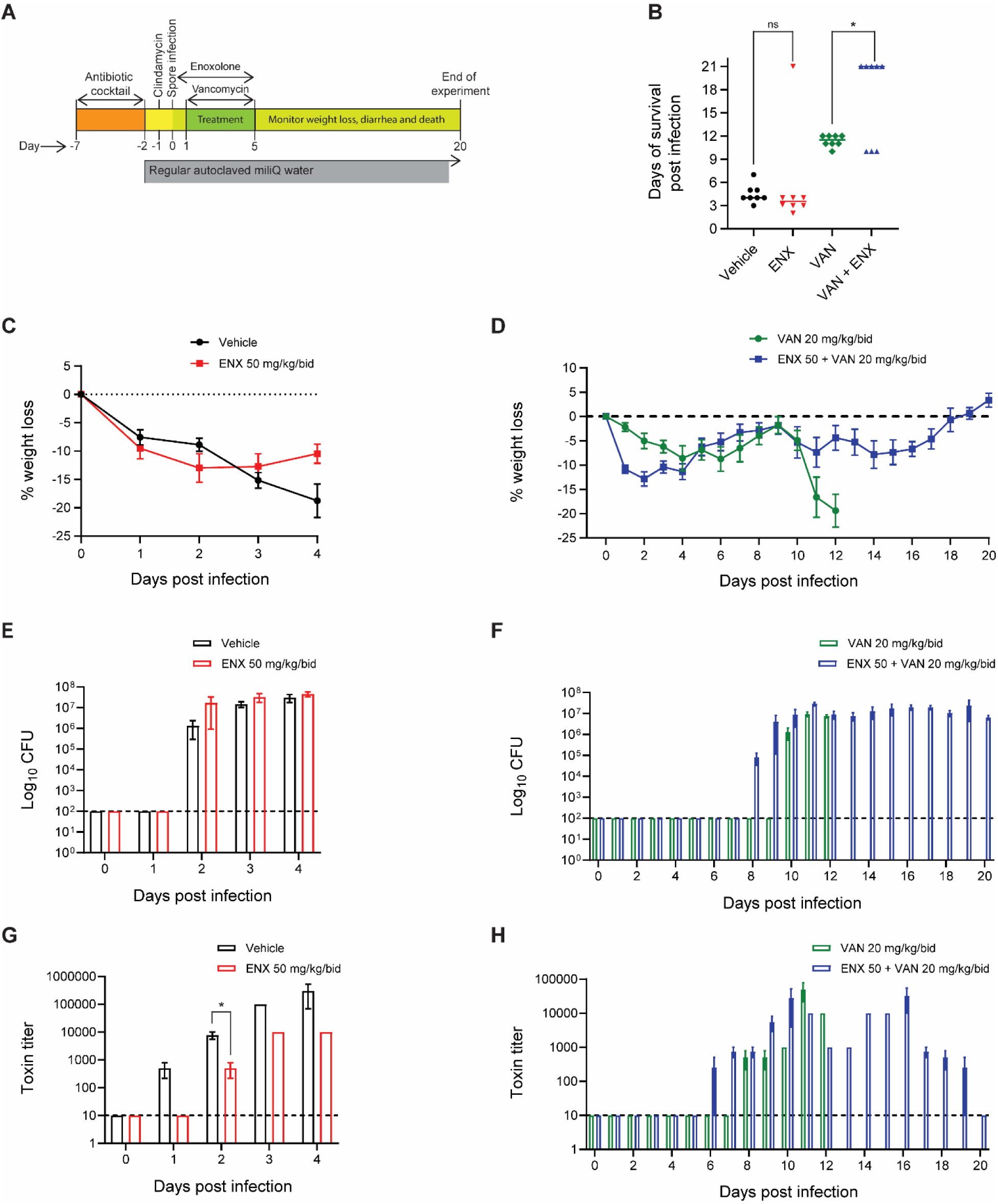
Efficacy of enoxolone (ENX) alone or in combination with vancomycin in mice with CDI colitis. (**A**) Schematic representation of the CDI colitis model. There were four groups of mice (4 males and 4 females in each group) as follows: vehicle (10% DMSO in corn oil), enoxolone (50 mg/kg), vancomycin (20 mg/kg) and combination (enoxolone 50 mg/kg, and vancomycin 20 mg/kg). Mice were treated twice daily (bid) via oral gavage. (**B**) Survival of mice following infection. Statistical analysis performed by unpaired t-test with Welch’s correction were as follows: ns (non-significant) for vehicle vs ENX; P<0.0001 for vehicle vs VAN; P<0.001 for vehicle vs ENX + VAN; P<0.05 for ENX vs VAN; P<0.01 for ENX vs ENX + VAN; and P<0.05, indicated as *, for VAN vs ENX + VAN. (**C and D**) Percent weight change, with respect to uninfected mice on day 0; shown are mice treated with vehicle or enoxolone in **C**; and mice treated with vancomycin or combination of enoxolone + vancomycin in **D**. (**E and F**) Analysis of *in vivo C. difficile* colonization (bioburden in feces) of mice treated with vehicle or enoxolone in **E**; and vancomycin or enoxolone + vancomycin combination in **F**; the limit of detection was 2 log_10_ CFU/g. (**G and H**) Toxin titers in feces, as determined by cell rounding against Vero cells, using feces from mice treated with vehicle or enoxolone in **G**; and vancomycin or enoxolone + vancomycin combination in **H**; the dashed line indicates limit of detection of 10^1^ toxin titer. In panels C, E and G animal numbers for vehicle and enoxolone treatment groups at days 4 were 7 and 4 mice, respectively.

### Efficacy of enoxolone in combination with vancomycin

Infected mice treated with vancomycin began to show signs of recurrence after treatment ended, and all 8 mice given vancomycin monotherapy succumbed by day 12 (**Figure 3B**). Combinatorial treatment with enoxolone and vancomycin reduced CDI development, as 62.5% (5 of 8 animals) of mice survived through to the experimental endpoint, day 20 (**Figure 3B** and **3D**). However, there were no substantial differences in the spore and toxin bioburdens between the vancomycin and combinatorial treatment groups (**Figure 3F** and **3H**). It is plausible that the enhanced efficacy of the combination may be due to known anti-inflammatory properties of enoxolone, including inhibition of TNF-alpha and interleukin-6 synthesis (27, 28), which could potentially reduce inflammation in CDI colitis. Further studies will be required to delineate the effects of enoxolone or related anti-inflammatory triterpenoid compounds on inflammation in CDI.

### Exploration of cellular targets of enoxolone

We next sought to explore the mechanism of action of enoxolone. To identify the cellular target(s) of enoxolone, we first employed click-chemistry combined with biotin-streptavidin pull down (**Figure S2A**). Hence, we synthesized an alkyne derivative of enoxolone (compound 3511) (**Figure S2B**), which we showed also inhibited toxin biosynthesis in R20291 (**Figure S2C**). Compound 3511 was immobilized to carboxymethylrhodamine (TAMRA) azide through CuAAC click reaction and then incubated with R20291 cell lysates. In parallel, a control mock reaction was performed with the native enoxolone. Visualization of click labeled proteins by in-gel fluorescence showed that bands of ∼50 to ∼60 kDa were prominent (**Figure S2D**) and suggested that enoxolone bound to multiple targets. Next, MALDI-TOF mass spectrometry was performed on proteins recovered on streptavidin magnetic beads. This identified 24 and 17 proteins in the click and mock reactions, respectively; of these, we eliminated 10 common proteins as background, leaving 14 proteins that were unique to the click labeled sample (**Table S2 and Figure S2E**). Six proteins were not considered further since their molecular weights were either <45 kDa or >70 kDa (outside of the molecular weight range seen from in-gel fluorescence). Of the remaining 8 proteins, we focused on GapN (NADP-dependent glyceraldehyde-3-phosphate dehydrogenase), AtpA (ATP synthase subunit alpha) and Ade (adenine deaminase), as their activities are predicted to affect metabolic networks that regulates toxin production. GapN is part of the glycolytic pathway that catalyzes the breakdown of glucose (29); ATP synthase is thought to couple ATP synthesis from the sodium/proton gradient generated from the coupled oxidation of ferredoxin and reduction of NAD+ by the RNF complex (29); and Ade catalyzes the early steps of the purine salvage pathway, in which it converts adenine to hypoxanthine that feeds into multiple pathways for purine biosynthesis (30). To first analyze GapN, AtpA or Ade effect on toxin biosynthesis, we quantified changes in toxin production following gene silencing. Antisense (asRNA) nucleic acid fragments were expressed from the Ptet promoter in pMSPT and induced by a growth permissive concentration of 0.032 µg/ml anhydrotetracycline (ATc) (**Figure S3A**) (24). Induction of asRNA for both *gapN* and *ade* caused moderate effects on growth while *atpA* showed no effects (**Figure 4A**). While transcripts for *atpA*, *ade* and *gapN* were reduced by ∼10 - 30 – fold (**Figure S3B**), the silencing of *atpA* and *ade* diminished TcdA production and transcription of *tcdA* and *tcdB* (**Figure 4B, 4C & S3C**). TcdA levels were reduced by 44.90 ± 17.35% and 68.71 ± 28.68% for *atpA* and *ade*, respectively. Ade and AtpA were therefore further studied, since the above findings suggest that inhibition of their activities affect toxin production and could be potential targets for enoxolone.

**Figure-4.**
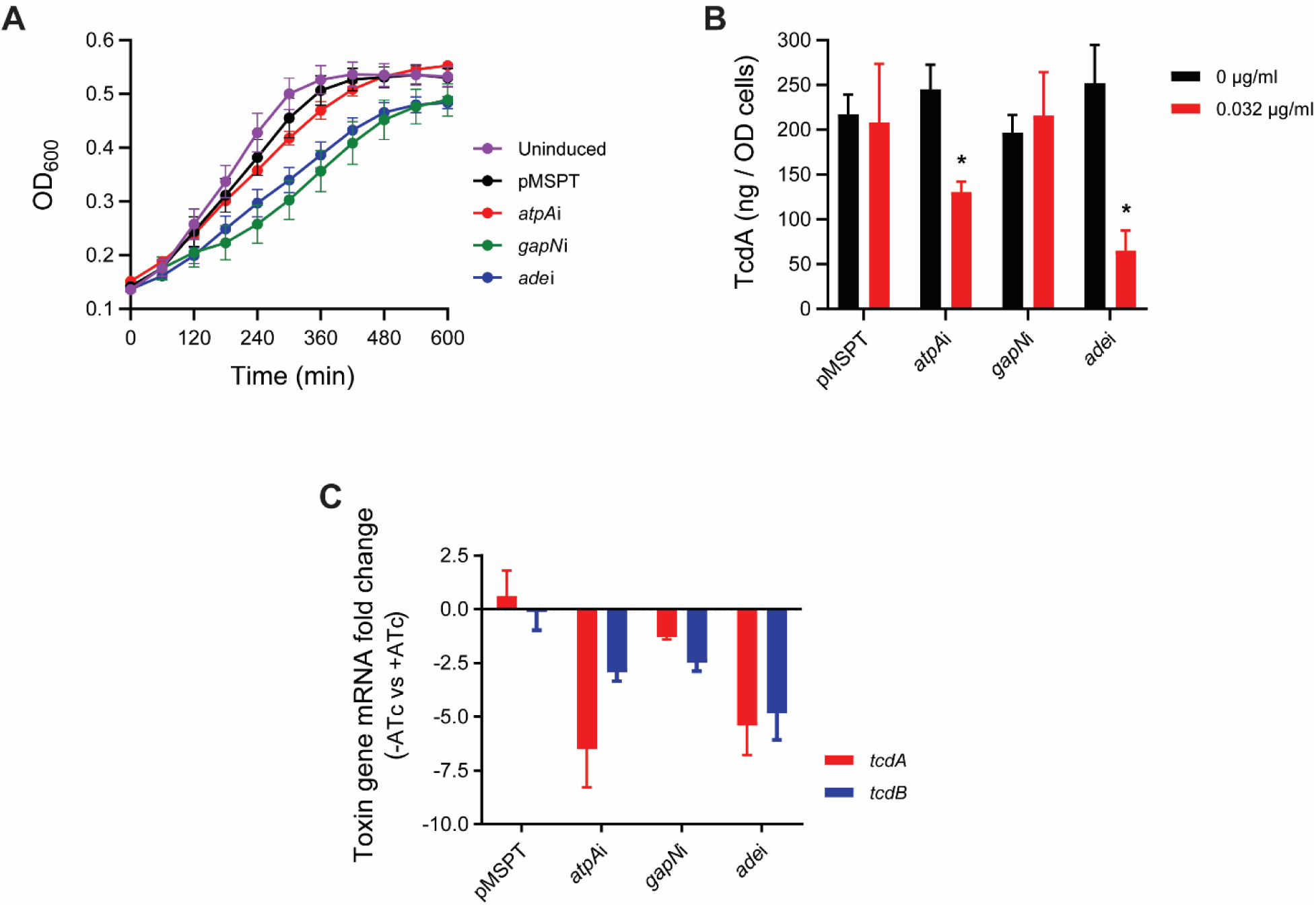
Effect of silencing genes for which proteomics suggested to encode targets of enoxolone. Analyzed were R20291 with the empty vector control (pMSPT) or R20291 with pMSPT expressing antisense RNA (asRNA) to *ade*, *gapN* or *atpA*; 0.032 µg/ml of anhydrotetracycline (ATc) was used for induction. **(A)** Growth kinetics were analyzed for the strains i.e., pMSPT (black), *atpAi* (red), *gapN*i (green) and *adei* (blue) in microdilution 96-well plate (n = 4 biological replicates). As a growth control, R20291 bearing pMSPT without ATc exposure was analyzed. **(B)** TcdA was analyzed at 24 h after induction of asRNA (n = 3 biological replicates). **(C)** mRNA levels of toxin genes (*tcdA* and *tcdB*) were analyzed by RT-qPCR (n = 3 biological replicates) from exponential cultures (OD_600_nm ≈ 0.3) treated with 0 or 0.032 µg/ml of ATc for 6 h. The fold change was calculated for mRNA from drug free and ATc exposed cultures.

### Enoxolone blocks the adenine deaminase activity and nucleotide metabolism

Surface Plasmon Resonance (SPR) was conducted to measure the binding of enoxolone to Ade. The Ade protein with N-terminal His-tag was adhered to the SPR chip via a His-tag antibody and testing of enoxolone at 5 to 400 µM indicated it bound to Ade in a concentration-dependent manner with a dissociation constant (K_d_) of 2.98 ± 3.2 µM (**Figure 5A**). The positive control 6-chloropurine, an Ade inhibitor (30), had a K_d_ of 25.17 ± 8.9 µM; the negative control vancomycin did not bind to Ade (**Table 2**). These observations were confirmed using Isothermal Titration Calorimetry (ITC), as the affinity values were 6 – 12 µM for enoxolone binding to Ade (**Figure S4**).

**Figure-5.**
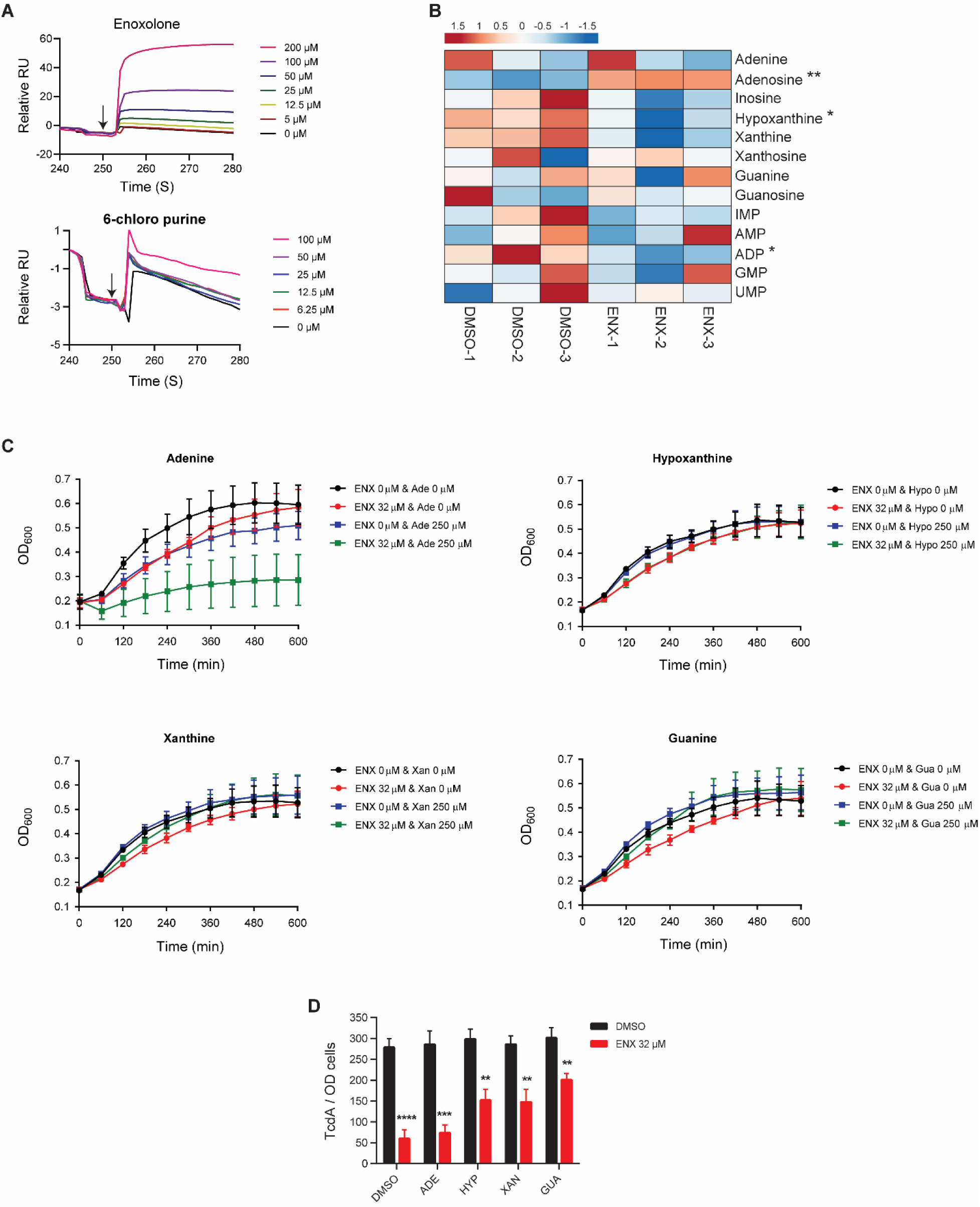
Enoxolone binds to adenine deaminase and inhibits purine metabolism. (**A**) Molecular interaction analysis using surface plasmon resonance (SPR). Dose response sensograms were attained with different concentrations of enoxolone (top panel) and 6-chloropurine (bottom panel) (shown in a representative of 3 replicates). **(B)** LC-MS/MS analysis of purine metabolites. *C. difficile* R20291 cells (OD_600_ ≈ 0.3) was exposed to enoxolone (16 µM) or 1% (v/v) DMSO for 3h. Heatmaps generated with Clustvis software shows relative quantities of metabolites; red and blue color intensities indicate levels of metabolites that were increased or decreased, respectively. **(C)** Growth kinetics of exponential R20291 (n = 3 biological replicates) in 1% (v/v) DMSO or enoxolone (32 µM) with or without various purine derivatives (250 µM). **(D)** Quantification of TcdA in R20291 cultures exposed to purine derivatives (250 µM) in presence of DMSO or enoxolone (32 µM); data is from 6 biological replicates and shown as means ± standard error of mean. Statistical analysis was performed by unpaired t-test with Welch’s correction **, P<0.01; ***, P<0.001; ****, P<0.0001 in Graphpad prism 9.3.1.

**Table-2.** Binding studies for enoxolone, 6-chloro purine and vancomycin with adenine deaminase protein by SPR.

| Compound | Kd (μM) | Rmax (RU) | Offset (RU) |
| --- | --- | --- | --- |
| Enoxolone | $2.98 \pm 3.27$ | $10.25 \pm 6$ | $-3.6 \pm 2.3$ |
| 6-chloropurine | $25.17 \pm 8.9$ | $3.41 \pm 2.45$ | $-0.29 \pm 2.23$ |
| vancomycin | ND* | $3.47^{+6} \pm 1.84^{+6}$ | $-3.6 \pm 2.3$ |

ND\* denotes not determined; RU= response units.

Because Ade participates in purine metabolism, converting adenine to hypoxanthine, we adopted targeted metabolomics to analyze intracellular changes to nucleotide metabolites. When R20291 cells were exposed to 16 µM enoxolone there were significant differential increases and decreases in the intracellular concentrations of adenosine and hypoxanthine, respectively (**Figure 5B**). Interestingly, ADP levels were significantly reduced in cells (**Figure 5B**). These results suggested that enoxolone disrupted purine metabolism, likely causing intracellular depletion of hypoxanthine. We therefore tested whether supplementation with purines affects the activity of enoxolone, in terms of its effects on growth and toxin production. Growth was inhibited when cells were exposed to adenine (250 μM) and this was more pronounced in the presence of enoxolone (**Figure 5C**). This suggests that intracellular accumulation of adenine is toxic to the cells. Hypoxanthine, xanthine, and guanine alone (at 250 µM) did not impact cell growth, but only xanthine and guanine marginally alleviated growth suppression by 32 µM enoxolone (**Figure 5C**). Toxin production was also partially restored by hypoxanthine, xanthine, and guanine, but adenine had no effect. Indeed, in the presence of 32 µM enoxolone, hypoxanthine or xanthine restored TcdA production by ∼30%. Greater (∼40%) recovery was seen with guanine (**Figure 5D**). Taken together, these observations indicate that adenine deaminase is blocked by enoxolone, which suppresses the downstream cellular pools of hypoxanthine, xanthine and guanine and their supplementation in media partially reverses enoxolone inhibition of toxin biosynthesis. Inability to fully restore toxin production also pointed to additional mechanisms, which were investigated below.

### Enoxolone depletes the cellular ATP levels

Attempts to purify AtpA to perform SPR failed (*data not shown*). We therefore next attempted target overexpression of *atpA* in R20291, under the Ptet promoter, to out-titrate enoxolone inhibition. However, this did not affect the EC_50_ of enoxolone (*data not shown*) and might reflect that overexpression of a single component of the F1F0-ATP synthase complex may not affect overall protein function. Hence, to determine if ATP synthase activity affects toxin biosynthesis, we tested Bz-423 a molecule that binds to the δ-subunit of F1F0-ATpase (31). At growth permissive concentrations of 8 and 16 µM, Bz-423 inhibited toxin biosynthesis (**Figure 6A**). These observations are in alignment with toxin biosynthesis being reduced upon *atpA* knockdown (**Figure 4C** **& S3C**) and support that ATP synthase activity affects *C. difficile* toxin biosynthesis.

**Figure-6.**
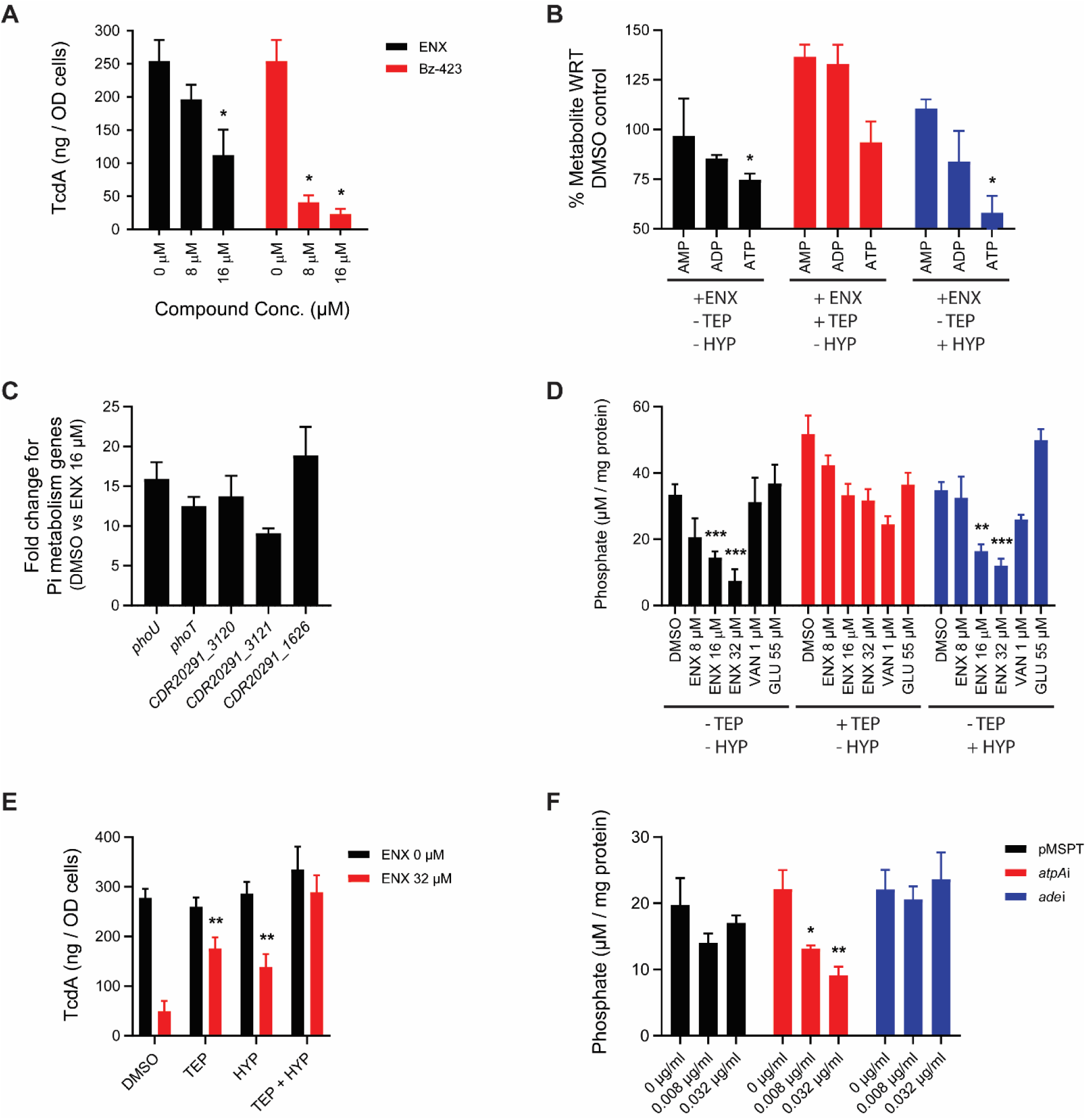
Role of ATP synthase activity and cellular phosphate pools on toxin production. **(A)** *C. difficile* R20291 (OD_600_ ≈ 0.3) was exposed to enoxolone (ENX, black bars) or Bz-423 (red bars). TcdA quantification was performed on culture supernatants after 24 of exposure. Data is representative of three biological replicates and shown as mean ± standard error mean (unpaired *t* test with Welch’s correction *, *P*<0.05 done using GraphPad prism version 9.3.1). **(B)** Quantification of intracellular pools of AMP, ADP, and ATP in *C. difficile* R20291 cells exposed to 16 µM of enoxolone. Cells in early exponential growth phase (OD_600_ ≈ 0.3) were exposed to DMSO or enoxolone in presence of 250 µM triethyl phosphate (red bars) or 250 µM hypoxanthine (blue bars). Cultures (n= 3 bioloical replicates) were harvested 3 h after exposure to enoxolone and metabolites were quantified through HPLC. The data in the plot is representative of three biological replicates and the error bars indicate means ± standard error of mean (SEM) and the data significance values were calculated to DMSO control (unpaired t-test with Welch’s correction *, P<0.05 in Graphpad prism 9.3.1). **(C)** Exponentially growing (OD_600_ ≈ 0.3, n=3 biological replicates) *C. difficile* R20291 cells were exposed to DMSO or 16 µM enoxolone for 30 min before extraction of RNA. mRNA levels for the genes involved in phosphate metabolism were analyzed by RT-qPCR, and the fold change was calculated as the difference in mRNA levels of control versus enoxolone-treated cells. **(D)** *C. difficile* cells at OD_600_ ≈ 0.3 were treated with 8, 16 or 32 µM of enoxolone, or 1 µM of vancomycin or 55 µM of glucose and the cells were harvested after 3 h. The phosphate content from respective cultures were analyzed form the whole cell lysates (n = 3 biological and 2 technical replicates). The significance of the data is calculated to DMSO control (unpaired t-test with Welch’s correction **, P<0.01; ***, P<0.001; ****, P<0.0001 in Graphpad prism 9.3.1). **(E)** TcdA quantification from 24 h old cultures of *C. difficile* R20291 cells exposed to 250 µM metabolite in presence of DMSO (black bars) or 32 µM enoxolone (red bars). The data in the plot is representative of minimum 3 biological replicates and the error bars indicate means ± SEM and the data significance is representative of unpaired t-test with Welch’s correction **, P<0.01 in Graphpad prism 9.3.1. **(F)** Relationship between AtpA activity and cellular phosphates, determined from phosphate levels in *C. difficile* R20291 cells carrying empty vector or the vector encoding an antisense RNA (asRNA) to *atpA* or *ade*. Cells were grown to OD_600_ ≈ 0.3 and asRNA induced with indicated concentrations of ATc. Cells were harvested after 3 h of exposure to ATc and their phosphate contents were measured from whole cell lysates (n = 3 biological and 2 technical replicates). The significance of the data is calculated for phosphate levels relative to uninduced cells (ATc 0 µg/ml) (unpaired t-test with Welch’s correction *, P<0.05; **, P<0.01 in Graphpad prism 9.3.1).

Since AtpA appeared to be a target, we tested if enoxolone affected cellular ATP levels. Here we quantified the nucleotide metabolites using HPLC in cells grown in the presence or absence of 16 µM enoxolone. The presence of enoxolone reduced cellular ATP by 25 ± 5.3% (**Figure 6B**). However, unlike observations from metabolomics, HPLC did not detect a significant depletion of ADP (**Figure 5B** **and 6B**). While these results do not directly show that enoxolone binds to AtpA, it establishes that enoxolone depletes cellular ATP in *C. difficile*.

### Enoxolone depletes cellular phosphates

Because enoxolone apparently has a multi-targeted action, we adopted RNAseq to identify its broader impact on cell physiology. Seventy genes were differentially expressed (FDR < 0.01 and fold change > 2.0) upon exposure to enoxolone (16 µM), of which 32 were downregulated and 38 were upregulated (**Figure S5A** and **Table S3**). RT-qPCR validation of the RNA-seq indicated a Pearson’s correlation of 0.8923 (p-value 0.0004) (**Figure S5B**). Pathway analyses revealed that approximately 42% of differentially expressed genes encoded membrane associated proteins, suggesting enoxolone imposed cell envelope stress; upregulated genes included cell wall biosynthesis (*CDR20291_2613*, *CDR20291_2614* and *CDR20291_2296*), antibiotic efflux transporters (*CDR20291_2117, CDR20291_2118, CDR20291_2296* and *CDR20291_2298*) and osmotic shock-related ion transporters (*CDR20291_1626, CDR20291_1718,* and *CDR20291_3147*). The most significantly upregulated genes involved phosphate metabolism genes, *CDR20291_1626* (a putative sodium/phosphate symporter) and genes of the Pho regulon (*CDR20291_3120* [encoding phosphate ABC transporter]*, CDR20291_3121* [encoding phosphate ABC transporter]*, phoT* [encoding PhoT phosphate import ABC transporter, ATP-binding protein] and *phoU* [encoding phosphate transport system regulatory protein PhoU] (**Table S3**). CDR20291_1626 is homologous to *Vibrio cholerae* NptA (32), a low affinity sodium dependent phosphate symporter. CDR20291_3119, CDR20291_3120 and CDR20291_3121 are homologous to PstB, PstA and PstC components of the PstSCAB high affinity phosphate transporter; *CDR20291_3129*, which is not part of the operon, encodes a PstS homolog, which was upregulated by 1.6-fold in the RNAseq (**Table S3**). Upregulation of the Pho regulon was confirmed by RT-qPCR since transcription of *phoU* was increased by 15.89 ± 3.69 – fold (**Figure 6C**). Activation of the Pho regulon and the NptA homolog occurs in response to decreases in inorganic phosphate, suggesting that enoxolone depleted inorganic phosphate. To test this, we quantified the intracellular phosphate in cells exposed to various concentrations of enoxolone for 3 h. Results showed that enoxolone depleted intracellular phosphate in a dose-dependent manner (**Figure 6D**). While 8 µM of enoxolone did not substantially affect cellular phosphates, 16 and 32 µM of compound depleted phosphates by 54.6 ± 17.6% and 75.6 ± 29.5% respectively. For comparison, cellular phosphate pools were not affected by vancomycin and glucose (**Figure 6D**).

To determine whether cellular phosphates impacted toxin production, we supplemented the growth medium with the organophosphate triethyl phosphate (250 µM) as a source for inorganic phosphate. In the presence of enoxolone (32 µM), triethyl phosphate restored intracellular phosphate (**Figure 6D**) and caused ∼60% restoration of toxin production (**Figure 6E**). However, triethyl phosphate did not appear to increase toxin production in control cells that were not exposed to enoxolone. To test if adenine deaminase had any impact on cellular phosphate pools, we quantified the intracellular phosphate levels for cells exposed to hypoxanthine (250 µM), in the presence and absence of enoxolone. As shown in **Figure 6D**, hypoxanthine did not restore cellular phosphate levels.

We next analyzed the influence of cellular phosphate levels on the metabolic pools of ATP by HPLC, showing that triethyl phosphate supplementation restored ATP levels in the presence of enoxolone (**Figure 6B**). Conversely, hypoxanthine, which did not affect phosphate levels, also did not affect cellular ATP (**Figure 6B**). These observations suggest that changes to phosphate levels were independent of adenine deaminase activity; furthermore, the effects of enoxolone on purine and phosphate metabolism are likely to occur via independent mechanisms. Thus, when media was supplemented with both triethyl phosphate and hypoxanthine there was full restoration of toxin A in the presence of 32 µM enoxolone (**Figure 6E**) i.e., restoration of phosphate and purine metabolism are required to fully bypass enoxolone inhibition.

### Silencing *atpA* depleted cellular phosphates

Since our findings clearly show that enoxolone depleted cellular phosphates, we determined if AtpA activity influenced phosphate metabolism. Hence, we quantified changes in cellular phosphates in cells expressing antisenses to *atpA* or *ade* relative to the empty vector control. Results showed that silencing *atpA* mRNA led to depletion of cellular phosphates pools, in manner dependent on the concentration of ATc inducer (**Figure 6F**). ATc at 0.008 and 0.032 µg/ml reduced cellular phosphates by 36.33 ± 17.2% and 56.14 ±15.75%, respectively. In contrast, silencing the *ade* mRNA had no effect on the pool of cellular phosphates (**Figure 6F**). Overall, these results indicate that ATP synthase activity is essential for *C. difficile* to main phosphate homeostasis.

## Discussion

From screening of structurally diverse phytochemicals, we identified that the triterpenoid enoxolone (18β-glycyrrhetinic acid) inhibits toxin biosynthesis by perturbing multiple metabolic pathways in *C. difficile*. Enoxolone is an aglycone metabolite of glycyrrhizin, the main bioactive component of licorice root extracts (*Glycyrrhiza* [sweet root]) (22). Licorice has a long history of use as a traditional Chinese medicine and is widely consumed in dietary supplements, herbal tea, confectionaries, beverages and cosmetics in the USA, Middle East, Asia, and Europe (22). Glycyrrhizin is hydrolyzed by intestinal bacteria to 18β-glycyrrhetinic acid that may undergo hepatic transformation to glucoronidated and sulfated conjugates that enter entero-hepatic circulation prior to renal excretion. The pharmacological properties of glycyrrhizin include anti-inflammatory, antipruritic, antioxidant and anti-infective activities, and are primarily due to enoxolone and its metabolites. It has also been reported that enoxolone is efficacious against staphylococcal skin infections in mice (28). Long-term use or overconsumption of licorice can lead to pseudoaldosteronism, which is characterized by hypertension and hypokalemia, resulting from enoxolone inhibition of 11-β-hydroxysteroid dehydrogenase type II enzymes that convert cortisol to cortisone (22). Noteworthy, glycyrrhizin the precursor for enoxolone did not inhibit toxin production (**Figure S6**), suggesting that the glucuronic acid disaccharide moiety hindered activity. Considering that the pharmacological properties of enoxolone are known and its short-term use has good safety profiles, we investigated its potential to be re-purposed as a CDI therapy. In our CDI colitis model, enoxolone improved the clinical outcome of mice treated with vancomycin. Although enoxolone treated mice had lower toxin loads it is unclear the extent to which this improved vancomycin treatment success, since the toxin and bioburden loads were similar between vancomycin and combination treatment groups. Several studies report that enoxolone inhibits the production of inflammatory cytokines (e.g., TNF-alpha, GM-CSF, and interleukin-6 (22), which are associated with CDI) (25). Thus, we speculate that the anti-inflammatory properties of enoxolone could have improved the prognosis of animals when it was given in combination with vancomycin.

By adopting enoxolone as a chemical genetic probe we discovered pathways that affect *C. difficile* toxin production and could be potential drug targets. Against strain R20291, enoxolone inhibited TcdA and TcdB biosynthesis at an EC_50_ of 16 μM and exhibited partial impact on growth at higher concentrations (≥32 μM). Through multiomics, genetic and biochemical approaches enoxolone was found to impede purine, phosphate, and energy metabolisms. This suggest it has a multi-target action, which is a finding that is consistent with reports that triterpenoids interact with multiple cellular targets (33, 34). Activity-based proteomics combined with gene silencing revealed Ade and AtpA as two of the molecular targets of enoxolone. Ade converts adenine to hypoxanthine, which is eventually metabolized to guanine nucleotides either via a xanthine intermediate or inosine (35). Metabolomic studies confirmed Ade as a molecular target where enoxolone reduced hypoxanthine content within cells, and the biochemical techniques SPR and ITC corroborated that enoxolone binds to Ade. These findings implicate the purine salvage pathway, and likely the *de novo* pathway, as having potential targets for inhibitors of *C. difficile* toxin production. This is also evident in other bacterial pathogens; for example, the *de novo* and salvage pathways converge at steps catalyzed by GuaB (IMP dehydrogenase), GuaA (GMP synthase) or Gmk (guanylate kinase) and disruption of *guaB* and *guaA* is known to attenuate the virulence of *Yersinia pestis* (*36*), *Salmonella* spp. (37) and *Francisella tularensis* (38). Conversely, deletion of *deoD* (encoding purine nucleoside phosphorylase) did not appear to affect *in vitro* virulence of *Mycobacterium tuberculosis* (39); DeoD cleaves purine nucleosides to their respective nucleobases following intracellular uptake. The salvage pathway is widely believed to be adopted by bacteria to maintain the GTP pool in conditions of low energy availability, to bypass the more energy intensive *de novo* pathway. Given our discovery that this pathway appears to modulate *C. difficile* virulence and there being a lack of knowledge of purine biosynthesis in this pathogen, further studies are warranted.

The effect of Ade inhibition on growth can be explained by two likely scenarios. Firstly, cellular build-up of adenine might be toxic to *C. difficile* since adenine supplementation attenuated growth. It is also plausible that Ade inhibition affected the downstream synthesis of guanosine phosphates that in turn reduced bacterial growth through multiple possible mechanisms. Indeed, GTP exhaustion blocks cell division by preventing polymerization of FtsZ, a cell division protein (40) that is a major cytoskeletal protein for cytokinesis and is sensitive to changes in purine metabolism (41). Although ATP depletion also affects growth, TEP supplementation that restored ATP levels did not enhance growth as was evident with xanthine and guanine supplementation (**Figure S7**). Thus, inhibition of purine metabolism affected both growth and toxin biosynthesis, while exhaustion of ATP and phosphate appear to mainly affect toxin biosynthesis.

Toxin biosynthesis was also inhibited by the ATP synthase inhibitor Bz-423 (31) at concentrations that did not inhibit growth (**Figure 6A**). Considering data from antisense knockdown of *atpA*, enoxolone depletion of ATP, the action of Bz-423 and metabolite analysis, there is an apparent correlation between ATP production via ATP synthase and toxin biosynthesis in *C. difficile*. Indeed, ATP synthase might be a key player in energy metabolism and redox biology. Recent analysis of *C. difficile* metabolism indicate that the ATP synthase complex is upregulated in the transcriptome and proteome of toxin producing stationary phase cells (42). ATP synthase in *C. difficile* is thought to play a key role generating ATP from the sodium/proton gradient generated by the RNF complex (29). Reduction of toxin production in *C. difficile* was reported to result from dissipation of the membrane potential (43), and an imbalance of the NAD^+^/NADH ratio (44). Our findings thus add to the growing indication that proteins involved in energy metabolism could be targets for inhibitors of toxin production.

Enoxolone depleted cellular inorganic phosphate, which is essential for the synthesis of ATP and other nucleotides and cellular signaling molecules. Bacterial cells adapt to phosphate starvation by activating the PhoB/PhoR two-component regulatory system leading to upregulation of phosphate acquiring PstSCAB transporter (45). In addition to maintaining phosphate homeostasis, the Pho regulon affects bacterial pathogenesis by regulating virulent gene expression (46). This is evident in *E. coli*, where an ExPEC Pst mutant (47) or a UPEC Pho mutant (48) were less virulent in chicken infection and mouse urinary tract infection models, respectively. Though the exact mechanism connecting Pho regulon to virulence gene expression is not well defined, we showed, as can be expected, that phosphate levels influence cellular ATP levels, which affects *C. difficile* toxin production. The reduction in cellular phosphate could be the result of changes to the clostridial membrane potential due to inhibition of ATP synthase, leading to a cycle where decreased phosphate also diminishes ATP biosynthesis.

In conclusion, chemical genomics with enoxolone enabled us to identify inter-related biological pathways that affect toxin production in C. *difficile* as depicted in the model diagram in **Figure 7**. However, limitations of this study include we could not quantify cellular pools of guanosine phosphates (*data not shown*), and the possibility that the improved efficacy for enoxolone combined with vancomycin is related to anti-inflammatory properties of enoxolone. Despite these limitations, from a microbiological standpoint, the multi-target action of enoxolone extends emerging knowledge that disruption of energy metabolism affects *C. difficile* virulence, and points to potentially druggable anti-virulent targets in *C. difficile*.

**Figure-7.**
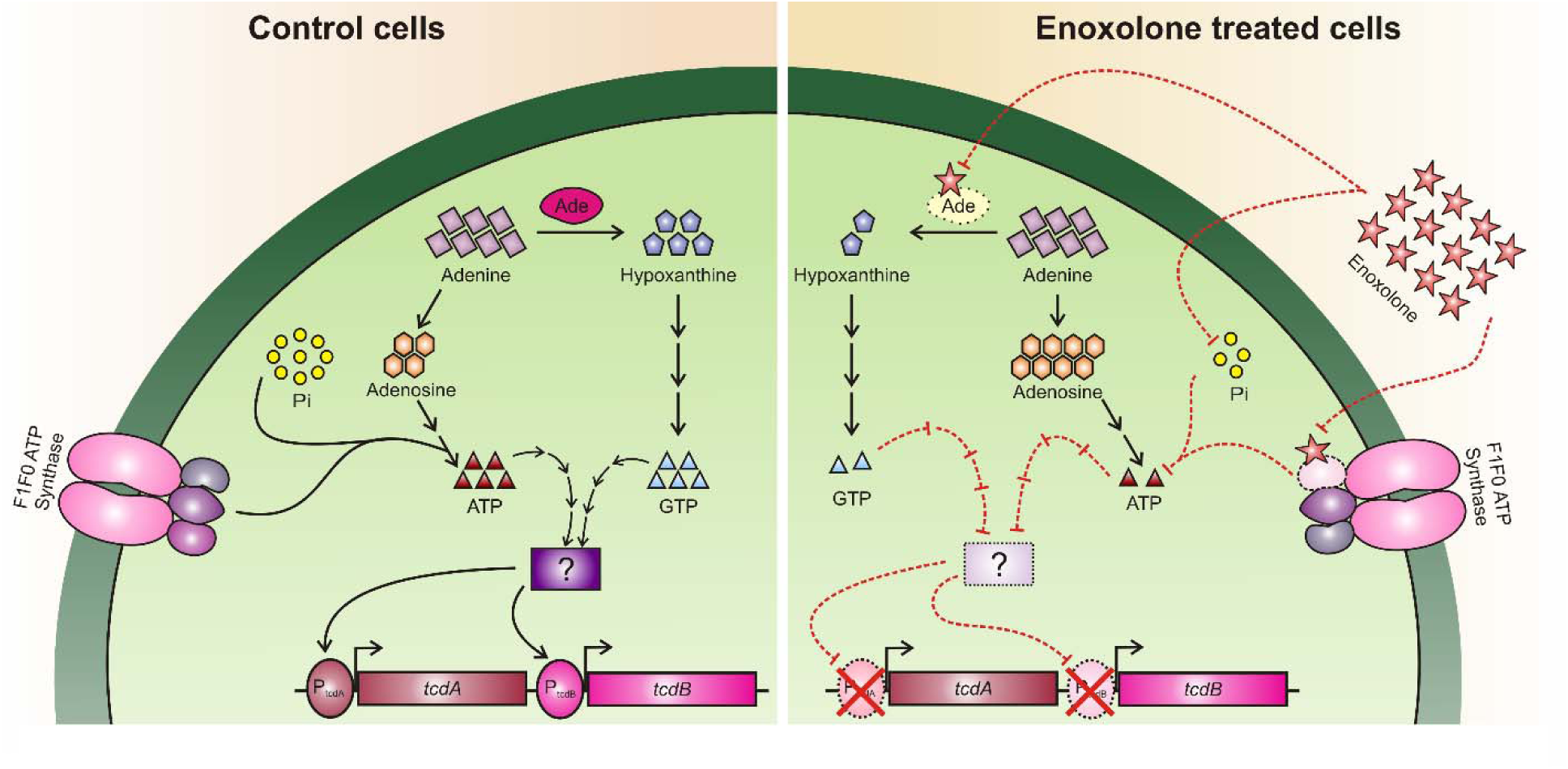
Proposed model for enoxolone mode of action against *C. difficile*. (Left) Scheme depicting the molecular events during the regular growth of *C. difficile*. (Right) represents adaptation of *C. difficile* to enoxolone, which hampers multiple processes in cells including ATP, phosphate, and purine metabolisms. The protein designated with a question mark, represents an unknown effector that might be a downstream factor causing repression of toxin genes. While effectors of toxin gene expression are known (e.g., TcdR, CcpA and CodY), it is not currently undetermined which regulator is modulated by enoxolone exposure.

## Materials and methods

### Bacteria and growth conditions

*C. difficile* strains were grown in Brain Heart Infusion (BHI) agar or broth and when needed supplemented with 0.1% (w/v) taurocholate or thiamphenicol (15 µg/ml). Gut flora species from Biodefense and Emerging Infectious Research Resource Repository (Manassas, VA) and the American Type Culture Collections (Manassas, VA) were grown in Brucella agar or broth supplemented with 5% (v/v) defibrinated sheep blood. Anaerobic cultures were grown at 37°C in anaerobic chamber (Don Whitley A35 anaerobic chamber). *E. coli* was grown aerobically at 37°C in Luria Bertani agar or broth with kanamycin (35 µg/ml), when appropriate.

### Susceptibility tests and growth kinetics

MICs were determined in BHI broth by two-fold serial dilution in 96-well microtiter plates. MICs were recorded as the lowest concentration of test compound that inhibited growth, following anaerobic incubation for 24 h. Growth kinetics in BHI broth were determined as described previously (44), using exponentially growing cells at OD_600_ ≈ 0.2. Automated growth was recorded every 60 min for 20 h with shaking before each read in an Infinite M Plex microplate reader (Tecan) in an anaerobic chamber (Coy Laboratory Products).

### Plasmid construction

Antisense fragments (100 bp), cloned into the anhydrotetracycline inducible vector pMSPT, were designed to target the ribosome binding site and start codon region as described (24). For overexpressing Ade protein, the codon optimized *ade* gene was cloned within the *Bam*HI/*Xho*I sites of pWL613a plasmid which is derived from pET28b by PPCMI core at Institute for Biosciences and Technology, Texas A&M.

### Toxin Quantification

Single colonies of *C. difficile* were inoculated into 5 ml of BHI broth. The resulting overnight culture was subcultured into fresh BHI medium at 1% v/v and grown to OD_600_ ≈ 0.2 to 0.3, before compounds or supplements were added to desired final concentrations. After 24 h of incubation cell densities (OD_600_nm) were recorded with Versa max microplate reader (Molecular Devices). After centrifugation, supernatants were collected and stored at -80°C for toxin quantification by cell rounding or ELISA, as described below.

#### (i) Cell rounding assay

Vero cells (50 µL) from ATCC were aliquoted in 384-well plates at a density of 10^4^ cells/ml and incubated overnight at 37°C in a CO_2_ incubator. Culture supernatants (100 nL) were added to the Vero cells with an Echo acoustics liquid handler (Labcyte) and the plates incubated at 37°C in a CO_2_ incubator for 3 h and then fixed with 4% (v/v) paraformaldehyde. After washing twice using a Hydrospeed washer (Tecan), cells were stained with 4′,6-diamidino-2-phenylindole (3.5 µM) for 15 min, washed and stained with 6 nM of Alexa Fluor 488 phalloidin stain (Invitrogen) for 2 h. Cells were then washed twice, before PBS (50 µL) was added. Images were recorded with an IN Cell Analyzer 6000 (GE Healthcare) for DAPI (excitation 405 nm and emission 455 nm) and Alexa Fluor 488 phalloidin (excitation 495 nm and emission 515 nm). Images were acquired from 4 fields/well at 10X magnification and exposure time of 1500 ms.

Image analysis was performed by calculating morphometric features using advanced imaging library in Pipeline pilot version 9.2 (Biovia). In brief, the background was corrected, followed by segmenting the nuclei and cell boundaries using thresholding and a watershed to separate neighboring objects. A panel of morphometric features were then extracted for both the nuclear and cellular mask. We trained a Random Forest classifier on a 50:50 randomly subsampling morphometric data from 16 toxin treated and 16 non-treated control wells, which were used as pseudo-labels for the cellular condition. Statistical evaluation of the model showed an area under the receiver operator characteristic curve for the out-of-bag training set to be 0.95. Gini analysis showed that that equivalent diameter, area, and convex area are the most important features used in classification. Confusion matrix analysis and other descriptive statistics of the with-held test set show a balanced accuracy of 0.93, indicating the model was robust. Given these observations, this model was applied to all morphometric analysis and reported as the proportion of rounded cells versus the total number of cells.

#### (ii) Toxin quantification by ELISA

TcdA and TcdB were quantified with *C. difficile* toxin A or B ELISA kit (tgcBIOMICS), according to the manufacturer’s instructions.

### Sporulation assay

This assay was performed as described previously (24). Briefly, cultures were grown anaerobically until OD_600_ ≈ 0.3, and exposed to compound for 5 days before enumerating total viable counts and spores.

### CDI colitis mouse model

Studies were conducted under an animal use protocol approved by Texas A&M University. The CDI colitis mouse model was adapted from a previously reported model (25), by using a lower concentration of dextran sodium sulphate (DSS, 1% w/v instead of 3% w/v) and administering an antibiotic cocktail and DSS for 5 days instead of 3 days. Antibiotics were kanamycin (0.4 mg/ml), metronidazole (0.215 mg/ml), vancomycin (0.045 mg/ml), gentamicin (0.035 mg/ml) and colistin (850 U/ml) (49). C57BL/6 mice (6 weeks), from Envigo, were treated with the antibiotic cocktail and DSS with 1% (w/v) for 5 days and 2 days later mice were given clindamycin (10 mg/kg via i.p.). About 20 h later, mice were challenged with gavaged 10^5^ viable spores of R20291. Oral gavage of enoxolone (50 mg/kg) or vehicle (10% DMSO in corn oil) was started ∼2 h after infection, whereas vancomycin (20 mg/kg) treatment was started at 24 h; mice were dosed twice daily. The dose of enoxolone was below its oral LD50 of >610 mg/kg in mice, but above the average daily intake of 100 mg for glycyrrhizin in humans, as recommended by the European Union (50). Mice were monitored thrice daily after infection for signs of morbidity (decreases in weight and temperature, and changes in appearance [wet tail, hunched posture, ruffled coat) and those that became moribund were euthanized. During the experiment, fecal samples were collected and stored at -80°C until use. CFUs were enumerated per gram of fecal pellet in 1 ml of sterile PBS. After heating at 65°C for 30 min, serial dilutions plated on selective and differential media for *C. difficile* (cycloserine-cefoxitin fructose agar with 0.1% (w/v) taurocholate). Toxin tires were quantified from the fecal pellets by cytopathic cell rounding assay.

### Synthesis of enoxolone click molecule, compound 3511

Compound 3511 was synthesized by mixing 1 mmol of enoxolone with 1.1 mmol each of propargylamine and 4-(4,6-dimethoxy-1,3,5-triazin-2-yl)-4-methylmorpholinium chloride (DMT-MM) in 10 mL methanol by stirring at room temperature for 5 h. After evaporation of methanol the residue was dissolved in 10 mL dichloromethane and sequentially extracted with 10 mL each of saturated aqueous NaHCO_3_, water, 5% citric acid, water, brine, dried over Na_2_SO_4_, and concentrated. The residue was purified by column chromatography using Hexanes/Ethyl acetate (3:1 - 1:1) to provide 241 mg (42%) of product as white solid. ^1^H NMR (400 MHz, CDCl_3_) δ 5.75 (t, *J* = 5.1 Hz, 1H), 5.62 (s, 1H), 4.10 – 3.90 (m, 2H), 3.16 (dd, *J* = 10.6, 5.7 Hz, 1H), 2.72 (dt, *J* = 13.5, 3.6 Hz, 1H), 2.27 (s, 1H), 2.18 (t, *J* = 2.5 Hz, 1H), 2.15 – 2.05 (m, 1H), 2.03 – 1.91 (m, 1H), 1.90 – 1.46 (m, 10H), 1.45 – 1.29 (m, 8H), 1.16 – 1.09 (m, 1H), 1.09 – 1.04 (m, 8H), 1.02 – 0.82 (m, 5H), 0.74 (d, *J* = 5.5 Hz, 6H), 0.63 (dd, *J* = 11.8, 1.9 Hz, 1H). ^13^C NMR (101 MHz, CDCl_3_) δ 200.15, 175.49, 169.02, 128.55, 79.74, 78.79, 71.70, 61.84, 54.97, 48.04, 45.37, 43.54, 43.21, 41.83, 39.18, 39.14, 37.36, 37.11, 32.78, 31.91, 31.46, 29.33, 29.30, 28.40, 28.11, 27.30, 26.48, 26.41, 23.36, 18.70, 17.49, 16.37, 15.57. MS-ESI, m/z = 508.38 [M+H]^+^.

### Click chemistry, affinity enrichment and protein identification

Overnight cultures of R20291 were diluted 1:100 into fresh BHI broth and grown to OD_600_ ≈ 0.5. Cells were harvested by centrifugation at 4000 x g for 15 min. The cell pellet was resuspended in buffer A (50 mM potassium phosphate pH 7.4, 150 mM NaCl) and supplemented with 10 mM MgCl_2_, 100 µg/ml of DNaseI and One cOmplete Protease Inhibitor (Roche). After cell lysis with a cell disruptor (Constant Systems Ltd), the lysates were centrifuged (10,000 x g for 15 min at 4°C). Membrane vesicles were isolated by centrifugation (100,200 x g for 25 min at 4°C) and the supernatant (cytosolic fraction) was collected. The membrane pellet was resuspended in buffer A with 10% (v/v) glycerol. Both the cytosolic and the membrane protein samples were aliquoted and stored at -80 °C until use. Protein concentrations were quantified using bicinchoninic acid (BCA) protein assay (Thermo Scientific), with bovine serum albumin as the calibration standard.

Click reaction between compound 3511 and TAMRA-biotin-azide (Click Chemistry tools) was performed in presence of 100 µM CuSO4, 500 µM tris-hydroxypropyltriazolylmethylamine and 5 mM Na-ascorbate. After incubation for 2 h in the dark at room temperature, the reaction was supplemented with cytosolic or membrane fractions (1 mg/ml) and incubated further for 4 h at 4°C. The clicked protein samples were then analyzed on 4 – 20% tris-glycine gradient gels (NuSep) and the TAMRA-biotin-azide conjugated protein samples were visualized by in-gel fluorescence using FluorChem M system (Protein Simple), with excitation 550 nm and emission 570 nm.

For the affinity pulldowns, 100 µl of streptavidin magnetic beads (Roche) were washed thrice with 10 volumes of buffer A. Protein samples were added to the beads and incubated overnight at 4°C on a rotating shaker (Barnstead Thermolyne). Next the beads were washed five times with 10 volumes of buffer A and bound proteins eluted with 0.1 M glycine-HCl, pH 2.5. Entire eluted samples were subjected to LC-MS/MS at the Proteomics Service Center, University of Texas Health Science Center.

### Protein expression and purification

To purify Ade, codon optimized *ade* was cloned into pWL613a, a derivative of pET28b plasmid, by standard cloning techniques and transformed into *E. coli* expression strain Bl21 (DE3). The strain was grown in LB medium with 50 µg/ml of kanamycin at 37°C until the OD_600_ ≈ 0.8. Cultures were cooled to 20°C, and protein expression induced with 500 µM isopropyl-*β*-D-thiogalactopyranoside (IPTG) for 16 h. Cells were harvested, lysed and soluble cytosolic fractions were collected following cell lysis with a cell disruptor. Ade was purified from Ni-NTA agarose beads (Marvelgent Biosciences Inc.) following pre-equilibration with buffer A containing 10 mM imidazole, pH 8.0 for 1 h, at 4°C on a rotating shaker (Barnstead Thermolyne). Proteins were eluted from Ni-NTA columns with buffer A containing 300 mM imidazole, pH 8.0 and further purified by size-exclusion chromatography on a Superdex 200 column (GE Healthcare) equilibrated with buffer A.

### Surface Plasmon Resonance

The binding of enoxolone to adenine deaminase protein was analyzed using Biacore T200 SPR machine (GE Healthcare), using a carboxymethylated CM5 chip (Cytiva). The CM5 chip was activated with 180 µl mixture of 10 mM NHS (*N*-hydroxysuccinimide) and ∼39 mM EDC (1-Ethyl-3-(3-dimethylaminopropyl) carbodiimide hydrochloride) at a flowrate of 7 µl/min. Monoclonal anti-histidine antibody was immobilized onto chip by passing 50 µg/ml anti-His antibody in 10 mM sodium acetate, pH 4.5 at a flowrate of 7 µl/min for 7 min (7000-15,000 RU). The recombinant Ade protein (100 µg/ml) was injected for trapping onto the chip through the immobilized anti-His antibody, with a contact time of 90 s at a flow rate of 10μL/min. The binding experiments were performed by injecting test compounds in 20 mM Tris pH 7.4 with 150 mM NaCl, 1% (v/v) dimethyl sulfoxide (DMSO) and 0.1% (w/v) BSA at a constant temperature of 25 °C. The anti-His antibody was regenerated by rinsing with 10 mM glycine pH 1.5, with a contact time of 30 s at a flow rate of 30 µl/min. Solvent correction was done with a 7-point solvent correction with DMSO concentrations ranging from 0.5% to 2.0% (all other buffer conditions were kept constant). Affinity values were determined using Biacore T200 Evaluation Software.

### Analysis of cellular metabolite concentrations

Overnight cultures of *C. difficile* R20291 were diluted 1:100 into fresh BHI broth. Cultures were grown to OD_600_ ≈ 0.3 and compounds added to desired concentrations. After 3 h the cultures were supplemented with equal volumes of chilled methanol and incubated at -20°C for 5 min. Samples were harvested by centrifugation (4000 x g for 10 min), washed with chilled buffer of 20 mM tris, 100 mM NaCl, pH 8.0. Cell pellets were stored at -80°C until further processing.

### Metabolome analysis by mass spectrometry

Cells were processed as previously described (51) and extracted samples analyzed by HPLC coupled to Agilent 6495 Triple Quadrupole mass spectrometry, using ESI positive ionization and Single reaction monitoring (SRM) mode. Nucleotides were identified in ESI positive mode using Zorbax eclipse XDB C-18, 1.8-micron, 4.6 x 100 mm column. Mobile phases A and B were 0.1% formic acid in water and acetonitrile, respectively. The gradient used is as follows: 0 min-2% B; 6 min- 2% of B, 6.5 min-30 % B, 7 min- 90% of B, 12 min 95% of B, 13 min 2% of B followed by re-equilibration at end of the gradient 20 min to the initial starting condition 2% of B. Flow rate: 0.2 ml/min. The data was processed using Mass Hunter Quantitative software. The data was normalized with internal standard and log2-transformed per-sample, per-method basis. For every metabolite in the normalized dataset, two sample t-tests were conducted to compare expression levels between different groups. Differential metabolites were identified by adjusting the p-values for multiple testing at an FDR threshold of <0.25 and generated a heat map.

### HPLC quantification of adenosine nucleotides

Cells were resuspended in 400 µl of 0.1 M perchloric acid, followed by disruption by probe sonication for 1 min (pulses for 5 sec on 50% duty cycle using Q125 Sonicator (Qsonica)). Cell lysates were centrifuged (10,000 x g for 10 min) and supernatants treated with 2.5% (v/v) of 3.5 M potassium chromate. Samples were filtered through 0.45 µM polyvinylidene difluoride membranes and analyzed by HPLC for purine metabolites at HPLC Bioanalytical core at Emory university.

### RNA Extraction and Sequencing

R20291 cultures (OD_600_ ≈ 0.3) were supplemented with either DMSO or enoxolone (16 µM). After 30 min, one volume of RNAprotect bacterial reagent (Qiagen) was added followed by centrifugation (4000 x g for 10 min). Cells were resuspended in 700 µl of Qiazol lysis reagent (Qiagen) and lysed in a Fastprep cell disruptor (Qiagen) for 5 min at force 50. Total RNA was extracted using the RNeasy Mini Kit (Qiagen), according to manufacturer’s instructions. Integrity of RNA was estimated on an Agilent Bioanalyzer 2100 with all samples possessing an RNA Integrity Number (RIN) >7. Ribosomal RNA was depleted with NEBNext rRNA Depletion Kit (New England Biolabs). Libraries were prepared from ribosomal-depleted RNA with the TruSeq Stranded mRNA Library Prep Kit according to the manufacturer’s instructions (Illumina, PN 20020595). Libraries were analyzed for insert size distribution using the 2100 BioAnalyzer High Sensitivity kit (Agilent). Libraries were quantified using the Quant-iT PicoGreen ds DNA assay (ThermoFisher) and by low pass sequencing with a MiSeq nano kit (Illumina). Paired end 100 cycle sequencing was performed on a NovaSeq 6000 (Illumina) at the Hartwell Center at St. Jude Children’s Research Hospital.

### Transcriptome analysis

The resulting reads were saved in FASTQ format and data was processed through the High-Performance Computing Facility at St. Jude. Adapter content and quality trimming was performed using Cutadapt (52). The quality of the raw and trimmed reads was assessed using FastQC and MultiQC (53). The resulting processed reads were aligned to the reference *Clostridium difficile* R20291 genome (Genome accession: FN545816) using STAR (54) and aligned reads were counted using featureCounts (55). Differential gene expression analysis was analyzed using DESeq2 (56) in the R statistical computing environment with significant differentially expressed genes determined using an FDR cutoff of <0.01 and an absolute value of log2(FC) > 1. The raw and processed RNA-seq data associated with this study have been deposited to NCBI GEO under accession number GSE199109.

### Gene expression analysis by RT-qPCR

The quantification of transcripts for various genes were analyzed by RT-qPCR as described previously (57) with slight modifications. The RNA samples were prepared as indicated above and the cDNAs were synthesized using qScript cDNA supermix (Quanta Biosciences) and RT-qPCR done with qScript SYBR Green RT-qPCR master mix (Quanta biosciences) in Applied Biosystems ViiA7 RT-PCR system. Results were calculated using the 2^-ΔΔCt^ method and transcript levels normalized to 16S rRNA.

### Phosphate quantification

Overnight cultures of *C. difficile* strains were diluted by 20-fold into a fresh medium and grown until OD_600_ ≈ 0.3. compounds or metabolites were added to the cultures at indicated concentrations and the cultures were incubated for 3h. cells were harvested by centrifugation (4,000 x g for 10 min) followed by washing cells five times with milliQ water. Cell pellets were resuspended in 1 ml of ice cold milliQ water and the cells were lysed by FastPrep (Qiagen) for 5 min and the samples were centrifuged at 21,100 x g for 5 min. Phosphate content was measured using the Malachite Green phosphate assay kit (Sigma Aldrich) according to the manufacturer’s instructions.

## Acknowledgements

This work was funded by grant R01AI144459 from the National Institute of Allergy and Infectious Diseases (NIAD) at the National Institutes of Health (NIH). The funders had no role in the study design, data collection and interpretation of the findings, or in the writing and submission of the manuscript. We thank Dharmanand Ravirajan from Institute for Biosciences and Technology (IBT), Texas A&M Health Science Center (TAMHSC) for technical assistance with ITC experiments. We thank Mary Sobieski from Institute for Biosciences and Technology, Texas A&M for technical assistance with data collection for cell rounding assays. RNA sequencing was carried out at the Hartwell Center for Biotechnology at St Jude Children’s research hospital, supported by the National Cancer Institute of the National Institutes of Health award P30 CA021765. The metabolomic analysis was performed at Metabolomics core at Baylor College of Medicine. The HPLC analysis was performed at HPLC Bioanalytical core at Emory University. The protein overexpression constructs were prepared at Protein Production, Characterization and Molecular Interaction (PPCMI) core at IBT TAMHSC; we are grateful to Magnus Höök (Director of PPCMI) for pre-reading of this manuscript. Protein mass spectrometry identifications were performed at Clinical and Translational Proteomics Service Center at University of Texas Health science Center in Houston.

## Supplementary figures

**Figure S1: Enoxolone (ENX) inhibits toxin production in various *C. difficile* ribotype and strains. (A & B)** Enoxolone dose response curves for toxin levels against *C. difficile* ribotype (RT) 027 strains CD196 (A) and UK1 (B). After exposing the exponential (OD_600_ ≈ 0.3) cells to various concentrations of Enoxolone for 24 h, toxins were quantified by ELISA for TcdA (red) and TcdB (blue) and by cytopathic cell rounding (green; CR). The EC_50_ observed for different RT027 strains through various methods are: R20291 (TcdA 11.38 μM; TcdB 14.29 μM; CR 7.77 μM), CD196 (TcdA 13.01 μM; TcdB 14.37 μM; CR 10.66 μM) and UK1 (TcdA 20.23 μM; TcdB 19.68 μM; CR 14.81 μM). **(C & D)** Effect of enoxolone on TcdA (**C**) and TcdB (**D**) production in varying *C. difficile* ribotype strains i.e., NR49296 (RT-014), NR49300 (RT-020), NR49310 (RT-078) and NR49318 (RT-106). Cultures were exposed to enoxolone at 8, 16 and 32 µM (based on the adopted EC_50_ against R20291 of 16 µM, which did not substantially affect growth). Toxin levels are relative to DMSO control. Data was obtained from three biological replicates and are presented as means ± standard error of mean (unpaired *t* test with Welch’s correction, *, *P*<0.05; **, *P*<0.01; ***, *P*<0.001; done using GraphPad prism version 9.3.1).

**Figure S2: Identification of molecular target(s) for enoxolone (ENX). (A)** Schematic of the click chemistry pull-down for target identification. Compound 3511 was coupled to carboxymethylrhodamine (TAMRA) azide through CuAAC click reaction followed by incubation with cell lysates. Proteins bound to the streptavidin-conjugated TAMRA azide were immobilized on streptavidin magnetic beads and the enriched proteins were identified by mass spectrometry analysis. **(B)** Synthetic scheme of compound 3511. The amine group from propargylamine was coupled to enoxolone in presence of 4-(4,6-dimethoxy-1,3,5-triazin-2-yl)-4-methylmorpholinium chloride (DMT-MM) in methanol at room temperature. **(C)** Quantification of TcdA from culture supernatants of R20291 exposed to equivalent concentrations of enoxolone (red bars) or compound 3511 (black bars) for 24 h. Data was from three biological replicates and shown as mean ± SEM (unpaired *t* test with Welch’s correction *, *P*<0.05; **, *P*<0.01; done using GraphPad prism version 9.3.1). **(D)** In-gel fluorescence detection for proteins bound to TAMRA biotin azide after click reaction followed by affinity purification with streptavidin magnetic beads. **(E)** Venn diagram showing overlapping and unique proteins identified from enoxolone mock and click reactions. The Venn diagram was generated through bioinformatics tool at VIB / UGent Bioinformatics & Evolutionary Genomics.

**Figure S3: Characterization of antisense knockdown constructs. (A)** Growth kinetics for the knockdown strains analyzed in a 96-well plate. Growth was analyzed for *C. difficile* R20291 harboring respective plasmids in different concentration of ATc (µg/ml) i.e., 0 (black), 0.008 (red), 0.016 (green), 0.032 (blue) and purple (0.064). Data obtained from four biological replicates are presented. **(B)** mRNA levels for the respective gene knockdowns i.e., *atpA* (red bar), *gapN* (green bar) and *ade* (blue bar) were analyzed by RT-qPCR. Cells were grown to OD_600_ ≈ 0.3 and the antisense RNA expression was induced with 0.032 µg/ml of ATc for 1 h. Fold changes were calculated as the difference of mRNA between untreated cells and those exposed to ATc (0.032 µg/ml). The data was from three biological replicates. **(C)** TcdA protein in R20291 with respective antisense knockdowns i.e., pMSPT (black), *atpA*i (red), *gapN*i (green) and *ade*i (blue) were cultured in different concentrations of ATc. After 24 h, TcdA was quantified in the culture supernatants. Data was from three biological replicates and shown as mean ± SEM (unpaired *t* test with Welch’s correction *, *P*<0.05; **, *P*<0.01; done using GraphPad prism version 9.3.1).

**Figure S4: Isothermal Titration Calorimetry (ITC) binding studies.** Characterization of enoxolone and adenine binding to adenine deaminase by ITC. Binding isotherms for the interaction of enoxolone (left panel), adenine (middle panel) and vancomycin (right panel) were generated by titrating the compounds into an ITC cell containing adenine deaminase (0.1 mM). Top panel depicts the heat differences upon injection of compounds and the bottom panel represents the integrated heat absorbed for injections. Data were fitted to a one site binding model, and the binding affinities calculated using Microsoft Origin software were: **enoxolone** Kd = 14.04 ± 11.11 µM, ΔG = 4.3 Kcal mol^-1^ and ΔH = 19.2 ± 0.7 Kcal mol^-1^; **adenine** Kd = 0.83 ± 1.36 µM, ΔG = - 7.1 Kcal mol^-1^ and ΔH = 1.8 ± 0.1 Kcal mol^-1^; **vancomycin** interactions were not detectable.

**Figure S5: Transcriptional response for *C. difficile* R20291 to enoxolone.** Gene expression was analyzed by quantifying mRNA in R20291 exposed to 16 µM (1 x EC_50_) enoxolone. Exponentially growing cells (OD_600_ ≈ 0.3) were exposed to enoxolone for 1 h before RNA was extracted for RNA-seq. **(A)** Volcano plot indicates differences in gene expression and statistical significance of expression changes was generated using GraphPad Prism (9.2.0). The plot shows upregulated (blue) and downregulated (red) genes between cells exposed to DMSO and 16 µM enoxolone. **(B)** Pearson correlation plot for a panel of 20 genes (11 – upregulated and 9 – down regulated) representing expression fold changes between RNAseq and RT-qPCR. The log_2_ fold change vales represent the calculated difference in mRNA levels between the cells exposed to DMSO and 16 µM enoxolone.

**Figure S6: Glycyrrhizin does not inhibit toxin production. (A)** The structure of glycyrrhizin consists of a glucuronic acid dimer linked to enoxolone by a glycosidic bond. **(B)** *C. difficile* R20291 grown to exponential phase (OD_600_nm≈ 0.3 were exposed to 100 µM glycyrrhizin or enoxolone; toxin levels for TcdA (red bars) and TcdB (blue bars) in culture supernatants were quantified. Data obtained from three biological replicates and shown as mean ± standard error mean (unpaired *t* test with Welch’s correction **, *P*<0.01; ***, *P*<0.001; done using GraphPad prism version 9.3.1).

**Figure S7: Growth kinetics (n = 3) for *C. difficile* R20291 under varying conditions.** Triethyl phosphate (TEP) supplementation did not restore growth in enoxolone (ENX), while unlike guanine (GUA) when added to cells exposed to enoxolone. Cells were exposed to 1% (v/v) DMSO or 32 µM enoxolone in the presence or absence of 250 µM TEP and/or 250 µM hypoxanthine (HYP) or 250 µM guanine. Growth kinetics were analyzed in a 96-well plate by reading absorbance at 600 nm for every 1 h for a span of 10 hrs.

## Supplementary tables

**Table S1:** Screened phytochemicals/metabolites in this study.

**Table S2:** List of proteins identified through affinity-based proteomics.

**Table S3:** List of genes that were differentially expressed after growth in presence or absence of enoxolone.

